# Enhancement of Macrophage Function by the Antimicrobial Peptide Sublancin Protects Mice from Methicillin-Resistant *Staphylococcus aureus*

**DOI:** 10.1101/299305

**Authors:** Shuai Wang, Qianhong Ye, Ke Wang, Xiangfang Zeng, Shuo Huang, Haitao Yu, Qing Ge, Desheng Qi, Shiyan Qiao

## Abstract

Methicillin-resistant *Staphylococcus aureus* (MRSA) is the major pathogen responsible for community and hospital bacterial infections. Sublancin, a glocosylated antimicrobial peptide isolated from *Bacillus subtilis* 168, possesses anti-bacterial infective effects. In this study, we investigated the role and anti-infection mechanism of sublancin in a mouse model of MRSA-induced sublethal infection. Sublancin could modulate innate immunity by inducing the production of IL-1β, IL-6, TNF-α and nitric oxide, enhancing phagocytosis and MRSA-killing activity in both RAW264.7 cells and peritoneal macrophages. The enhanced macrophage function by the peptide *in vitro* correlated with stronger protective activity *in vivo* in the MRSA-invasive sublethal infection model. Macrophages activation by sublancin was found to be mediated through the TLR4 and the NF-κB and MAPK signaling pathways. Moreover, oral administration of sublancin increased the frequencies of CD4^+^ and CD8^+^ T cells in mesenteric lymph nodes. The protective activity of sublancin was associated with *in vivo* augmenting phagocytotic activity of peritoneal macrophages and partly improving T cell-mediated immunity. Macrophages thus represent a potentially pivotal and novel target for future development of innate defense regulator therapeutics againt *S. aureus* infection.

---

Concurrent with the success of antibiotics for treating infections, their excessive use contributes to the emergence of antibiotic-resistant bacteria (1). Methicillin-resistant *Staphylococcus aureus* (MRSA) is widespread and multi-resistant, thus has challenged the effectiveness of antibiotics including β-lactams, macrolides, and quinolones, as well as vancomycin which often is the last drug that successfully treats MRSA infections (2). Antibiotic resistance has become an increasingly serious health care problem in the world (3). This has been aggravated by a collapse in the number of approvals of new antibacterials in the past three decades (4).

Macrophages are professional phagocytes of the innate immune system, providing a first line of defense against infections. It has been reported that macrophages played an important role in the clearance of *S. aureus* in the infected mice (5). Mice that were depleted of macrophages are susceptible to MRSA infection (6). Nevertheless, some investigators have pointed out several characteristics that MRSA may thwart the macrophage-mediated host defense (7). Macrophages can kill bacteria directly through phagocytosis and indirectly via releasing inflammatory molecules, nitric oxide (NO), as well as by secreting pro-inflammatory factors, such as interleukin-6 (IL-6), IL-1β, and tumor necrosis factor-α (TNF-α) (8, 9). Macrophages are the first immune cells that are recruited to the infection site, and are the main source of pro-inflammatory cytokines after activation (10). Many investigators have reported progressing macrophage dyfunction may contribute to severe sepsis (11, 12).

Antimicrobial peptides (AMPs) are important components of the innate immune defense against a wide range of invading pathogens (13, 14). Sublancin is a 37 amino acid AMP isolated from the Gram-positive soil bacterium *Bacillus subtilis* 168 (15). In our previous studies, we showed that sublancin was protective in several *in vivo* infection models. Although the minimum inhibitory concentration (MIC) of sublancin was much higher than that of traditional antibiotics *in vitro*, it was demonstrated that sublancin was effective against *Clostridium perfringens* induced necrotic enteritis in broilers (16). We also found that sublancin has potential for prevention of *S. aureus* infection in mice (17). Moreover, sublancin was further found to protect against drug-resistant bacteria in a mouse MRSA infection model (18). Several reports have been demonstrated that AMPs were capable of activating macrophage function (13, 19). Recently we revealed the capability of sublancin in activating macrophages and improving the innate immunity of mice *in vivo* (Wang et al., accepted for publication). Hence, the goal of the present study was to explore the potential anti-infection mechanism of this peptide. In the present study, we investigated whether sublancin can (i) activate macrophages and the signaling pathway involved in this process, (ii) inhibit bacterial growth in a model of MRSA infected mice and macrophages, (iii) improve immune function in mice under healthy and MRSA-induced sublethal infection conditions.

## RESULT

### Sublancin promoted the secretion of cytokines and NO from RAW264.7 cells and mouse peritoneal macrophages

Activated macrophages produce cytokines such as IL-1, IL-6 and TNF-α and also secrete cytotoxic and inflammatory molecules such as nitric oxide (NO) (20, 21). Hence, the effect of sublancin on cytokine production by RAW264.7 cells and mouse peritoneal macrophages (P-Mac) was investigated. Untreated RAW264.7 cells secreted a basal level of TNF-α but barely detectable amounts of IL-1β and IL-6 (Fig. 1). The addition of sublancin significantly stimulated the production of IL-1β, IL-6 and TNF-α in a concentration dependent manner (*P* < 0.05). To study the effect of sublancin on NO production in RAW264.7 cells, we measured the secretion of NO in the culture supernatant of RAW264.7 cells stimulated with sublancin alone. Compared to the control, the addition of sublancin resulted in remarked increase in NO production in a dose-dependent manner (*P* < 0.001; Fig. 1). Similar results were observed in P-Mac.

**FIG 1.**
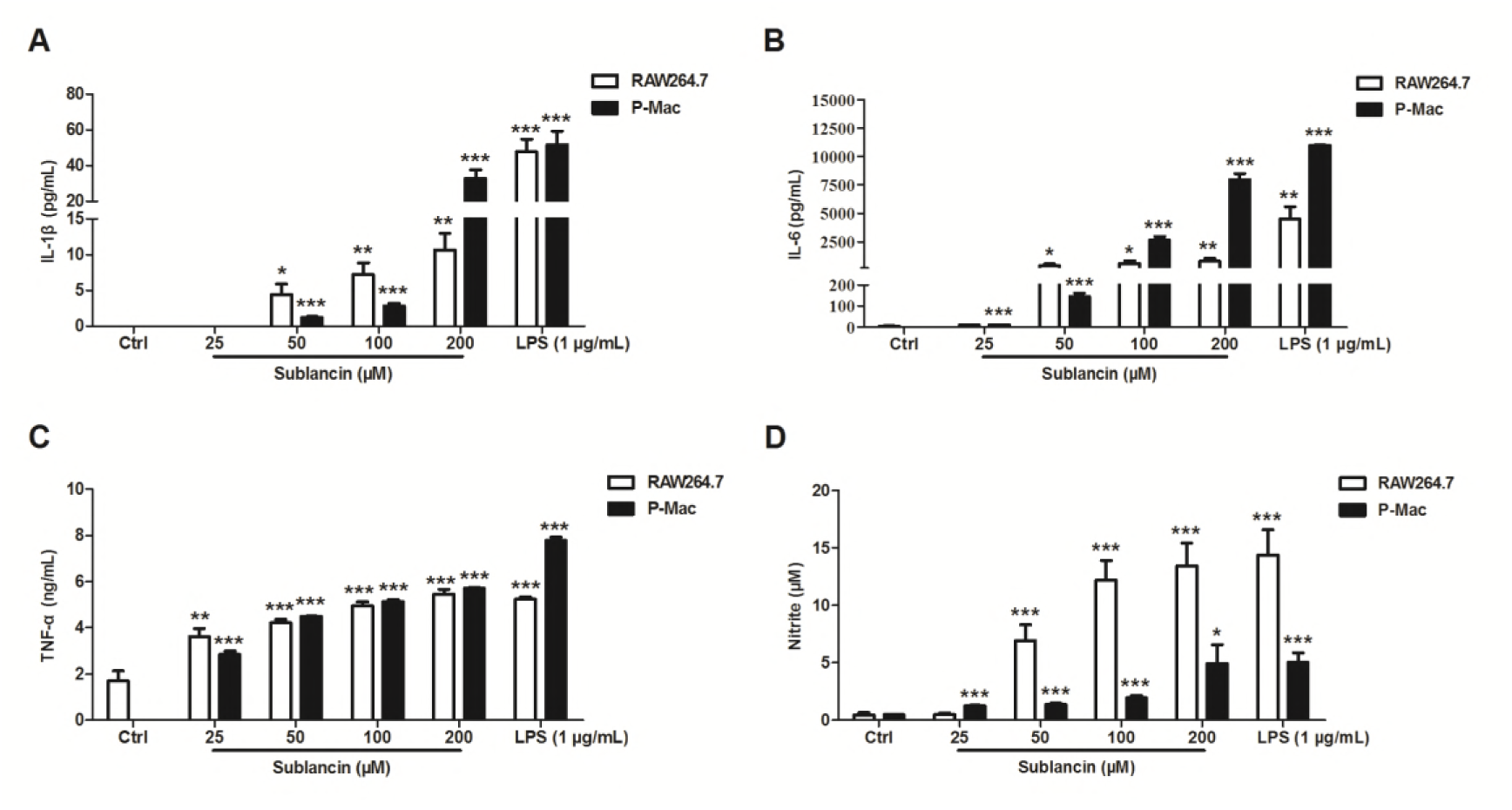
Effect of sublancin on the production of cytokines (A to C) and nitric oxide (D) from RAW264.7 cells and mouse peritoneal macrophages (P-Mac). RAW264.7 cells and P-Mac were treated with sublancin (0-200 μM) or LPS (1 μg/mL) for 24 h. The data are expressed as mean ± SEM (n = 6). Significant differences with control cells were designated as \**P* < 0.05, \*\**P* < 0.01 or \*\*\**P* < 0.001

### Sublancin regulated mRNA expression of inflammatory factors, chemokines and costimulatory molecules in RAW264.7 cells and peritoneal macrophages

Due to the crucial role of inflammatory factors, chemokines and costimulatory molecules in the activation and function of macrophages, we used quantitave RT-PCR to investigate the potentials for sublancin to regulate the expression of these mediators in RAW264.7 cells and P-Mac at the mRNA level. Consistent with the results at the protein level, a significant upregulation of IL-1β, IL-6, TNF-α and iNOS mRNA expression was seen in the RAW264.7 cells or P-Mac treated with sublancin (*P* < 0.05) (Fig. 2). COX-2 is an important upstream regulator for prostaglandin E_2_ expression. The sublancin treatments resulted in notable increase in the mRNA expression of COX-2 than the control cells. IL-8 and monocyte chemoattractant protein-1 (MCP-1) are the primary chemokines that recruit neutrophils and monocytes, respectively. We found that sublancin increased mRNA expression of IL-8 and MCP-1 in both RAW264.7 cells and P-Mac. B7-1 and B7-2 are two classical surface markers of activated macrophages. It was found that the expression of B7-1 and B7-2 was also increased significantly in sublancin treatments, suggesting that sublancin directly induces macrophage activation.

**FIG 2.**
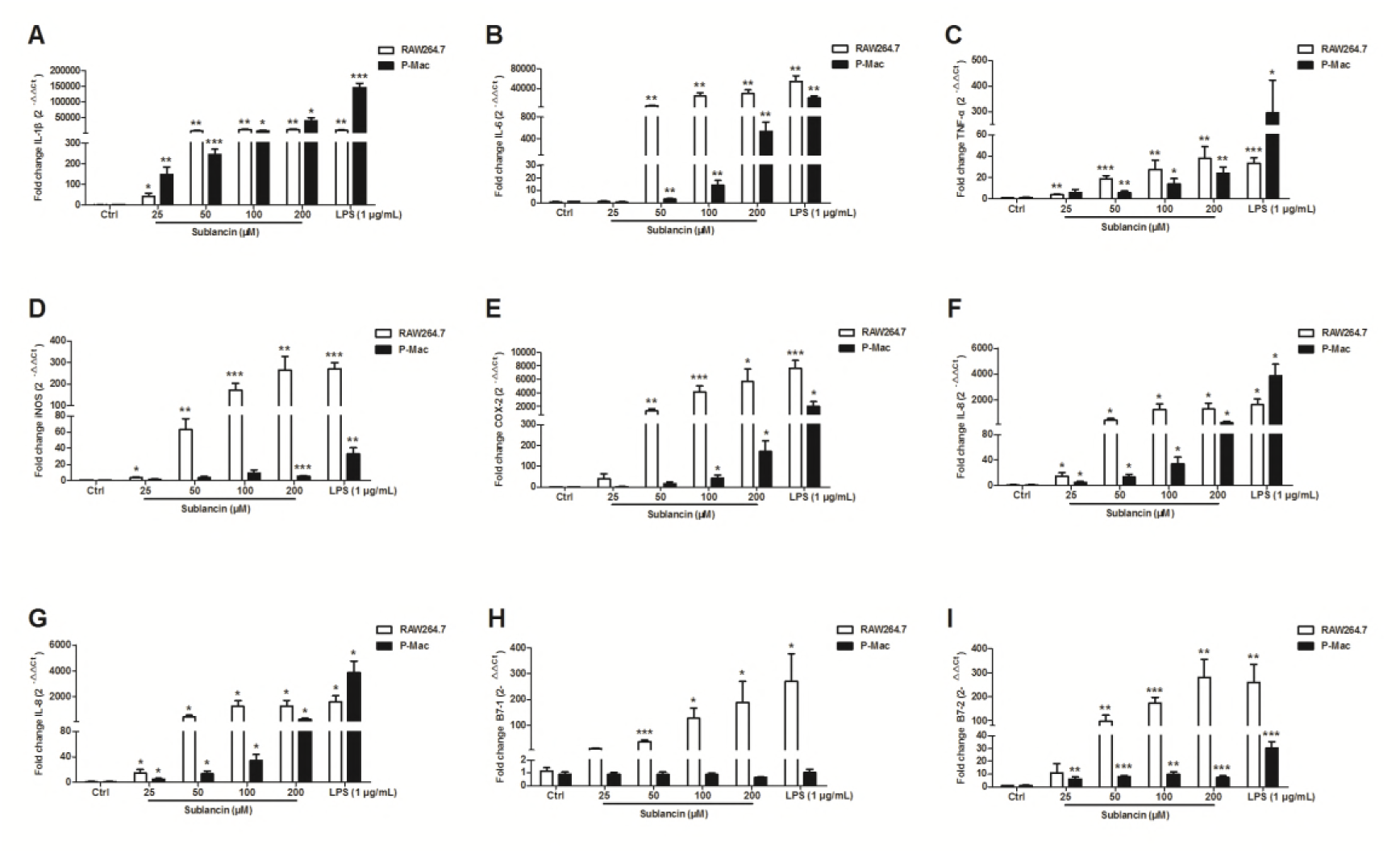
The mRNA expression of inflammatory factors (A to E), chemokines (F and G) and costimulatory molecules (H and I) in RAW264.7 cells and mouse peritoneal macrophages (P-Mac) treated with sublancin. RAW264.7 cells and P-Mac were treated with sublancin (0-200 μM) or LPS (1 μg/mL) for 12 h. The expression of target genes was detected by real-time PCR. GAPDH was used as an internal standard for normalization. The values are presented as mean ± SEM (n = 6). Significant differences with control cells were designated as \**P* < 0.05, \*\**P* < 0.01 or \*\*\**P* < 0.001

### Influence of sublancin on macrophage phagocytic activity

Phagocytosis is one of the primary functions of macrophages, and it is specialized in excluding foreign bodies (22). We examined the effect of sublancin on phagocytic uptake of fluorescent microspheres in RAW264.7 cells and peritoneal macrophages using flow cytometer. As shown in Fig. 3, sublancin stimulated the phagocytotic activity of RAW264.7 cells or peritoneal macrophages compared with the control.

**FIG 3.**
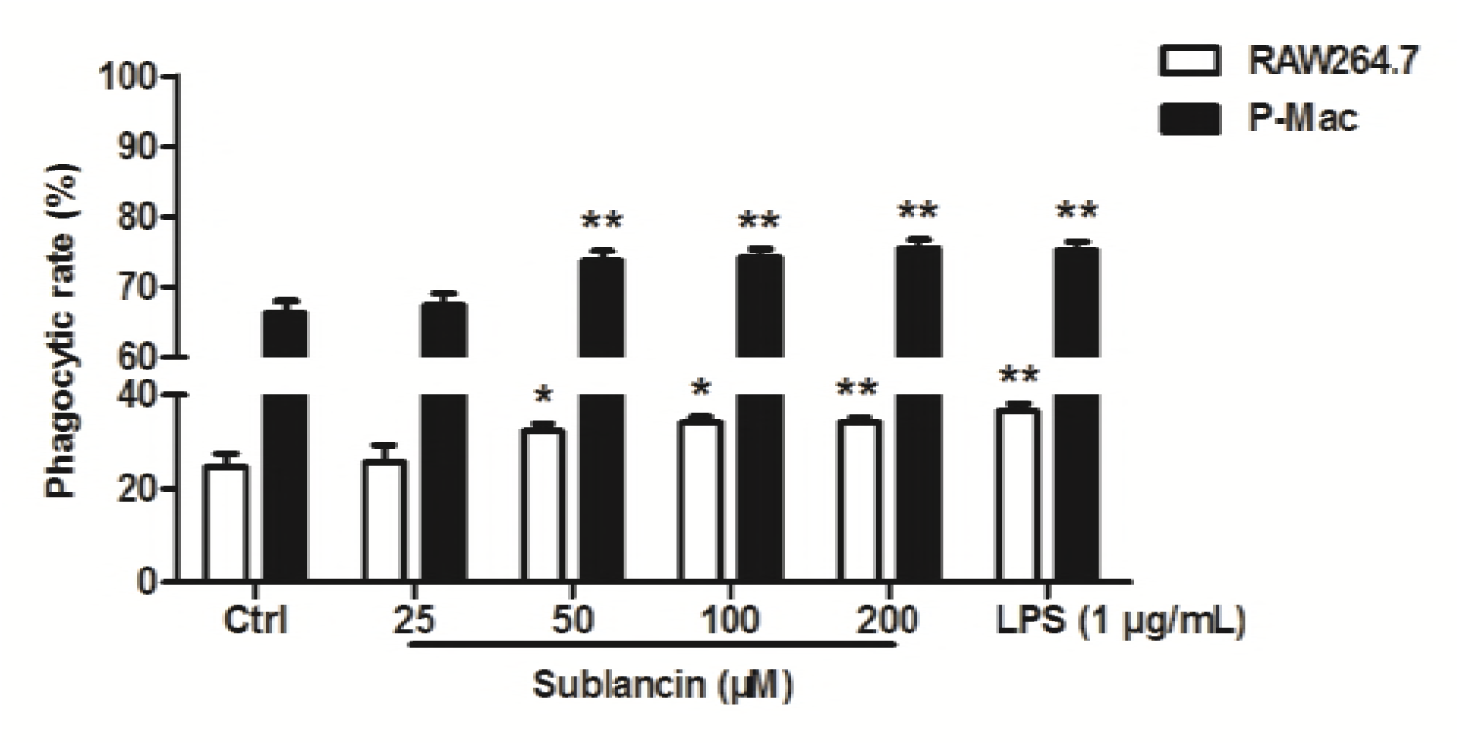
Effect of sublancin on phagocytic activity of RAW264.7 cells and mouse peritoneal macrophages (P-Mac) *in vitro*. RAW264.7 cells and P-Mac were treated with sublancin (0-200 μM) or LPS (1 μg/mL) for 12 h. Phagocytic activity of macrophage cells was assessed in terms of the population of phagocytic cells relative to the total number of cells for RAW264.7 cells and P-Mac. The values are presented as mean ± SEM (n = 6). Significant differences with control cells were designated as \**P* < 0.05 or \*\**P* < 0.01

### Sublancin enhances bactericidal capacity of macrophages

Phagocytosis is the first step of the bactericidal activity of macrophages. After 1 hour, the phagocytosis of MRSA remained unchanged in RAW264.7 cells or peritoneal macrophages preincubated with 25 μM sublancin. Bacterial killing by macrophages was assayed after 24 hours of incubation. We observed that a marked decline in viable bacteria occurred in sublancin treated cells (Fig. 4).

**FIG 4.**
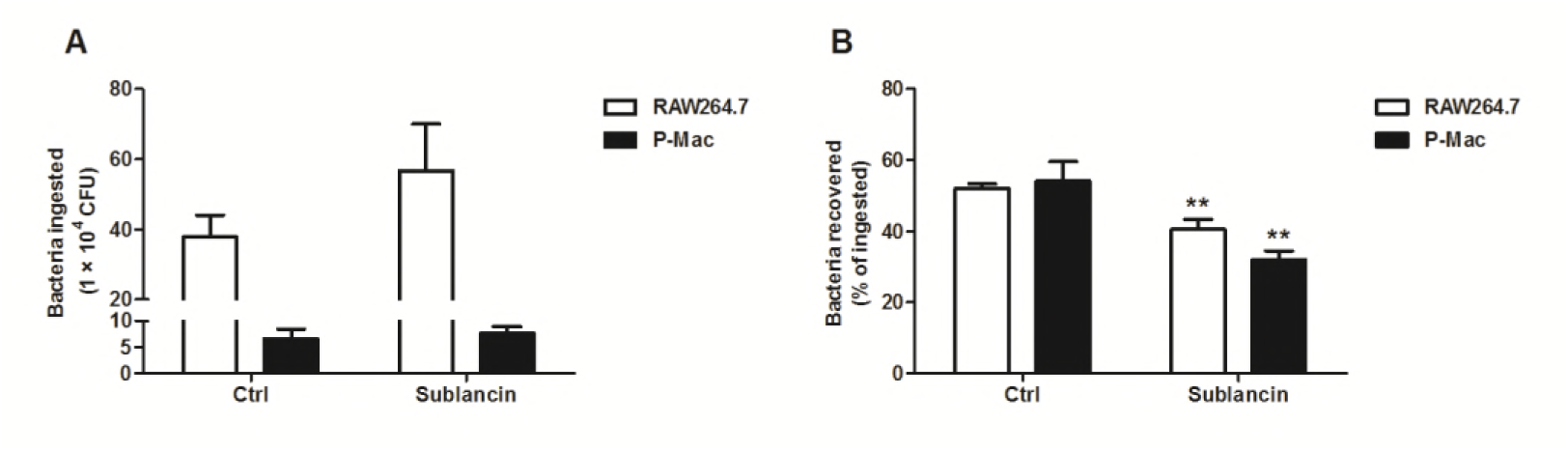
The antimicrobial peptide sublancin promoted the capacity of RAW264.7 macrophages and mouse peritoneal macrophages (P-Mac) to kill *Staphylococcus aureus* ATCC43300. RAW264.7 macrophages and P-Mac were cultured 12 hours with or without sublancin (25 μM) and then exposed to ATCC43300 (MOI 20). Phagocytosis (A) and killing (B) of bacteria were quantified as described in Materials and Methods. Data are presented as mean ± SEM (n = 6). Significant difference with control cells was designated as \*\**P* < 0.01

### Sublancin activates RAW264.7 cells through TLR4 signaling pathways

TLR4 plays a critical role in the activation of innate immune response by recognizing specific molecular patterns. To explore whether TLR4 is involved in sublancin-induced macrophage activation, RAW264.7 cells were pre-treated with TLR4 inhibitor (TAK-242). Then, the mRNA expression of cytokines were detected using quantitave RT-PCR. As shown in Fig. S1A to C, with the presence of TLR4 inhibitor, the mRNA expression of IL-1β, IL-6, and iNOS induced by sublancin was drastically suppressed and was significantly lower than those without inhibitor (*P* < 0.001). The results show that the immunostimulatory effect of sublancin in macrophages is exerted probably through activation of TLR4. The stimulation of TLR4 signaling pathway ultimately triggers the activation of phosphorylation of MAP Kinases and transcription of NF-κB (23). MAPKs are well conserved protein kinases including p38 MAP kinases, extracellular signalregulated kinase, and JNK (24-26). Treatment with 100 μM sublancin resulted in a significant increase in the phosphorylation of all three MAPKs (p38, ERK, and JNK) and the phosphorylation peaked 30 min after sublancin exposure (Fig. 5). We also investigated the phosphorylation status of the above mediators after treatment with sublancin at the different concentration for 30 min. Sublancin significantly upregulated the phosphorylation of all the three MAPKs in RAW264.7 cells in concentration-dependent manner (Fig. 6). In macrophages, NF-κB is an important regulator of immune activation through induction of many cytokines (27). Sublancin stimulated phosphorylation of NF-κB p65, while it correspondingly decreased IκB-α in the cytosol (Fig. 6). To further confirm whether sublancin activates macrophages through MAPK and NF-κB pathways, we used inhibitors to inhibit the initiation of signal transduction. Sublancin-induced mRNA expression of IL-1β and IL-6 was significantly suppressed by NF-κB (Bay11-7082) inhibitor (*P* < 0.05) (Fig. S1D, S1E). In addition, the ERK (U0126) and p38 (SB203580) inhibitors exerted significantly inhibition of iNOS mRNA expression (*P* < 0.001) (Fig. S1F). Moreover, the phosphorylation of p38 MAPK, ERK1/2, and JNK induced by sublancin was dramatically inhibited by TLR4 inhibitor (Fig. S1G). These data strongly indicated that the activation of macrophages induced by sublancin was mediated by TLR4 signaling pathways.

**FIG 5.**
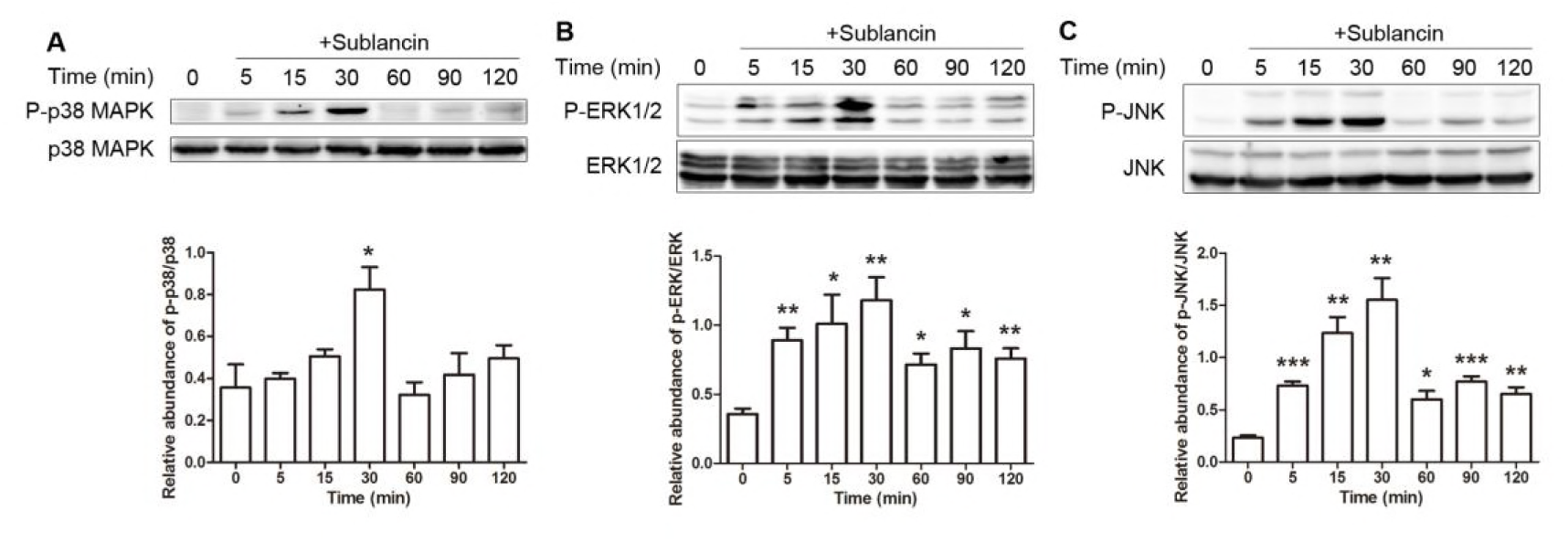
Analysis of p38, ERK, and JNK signaling pathways in RAW264.7 cells treated with sublancin for different times. RAW264.7 cells were treated with 100 μM sublancin for the indicated times and the phosphorylation of p38, ERK1/2, and JNK were detected by Western blot analysis. Representative immunoblots and quantitation of the phosphorylation abundance of p38 (A), ERK1/2 (B), and JNK (C). The values are presented as mean ± SEM (n = 3). Significant differences with control cells were designated as \**P* < 0.05, \*\**P* < 0.01 or \*\*\**P* < 0.001

**FIG 6.**
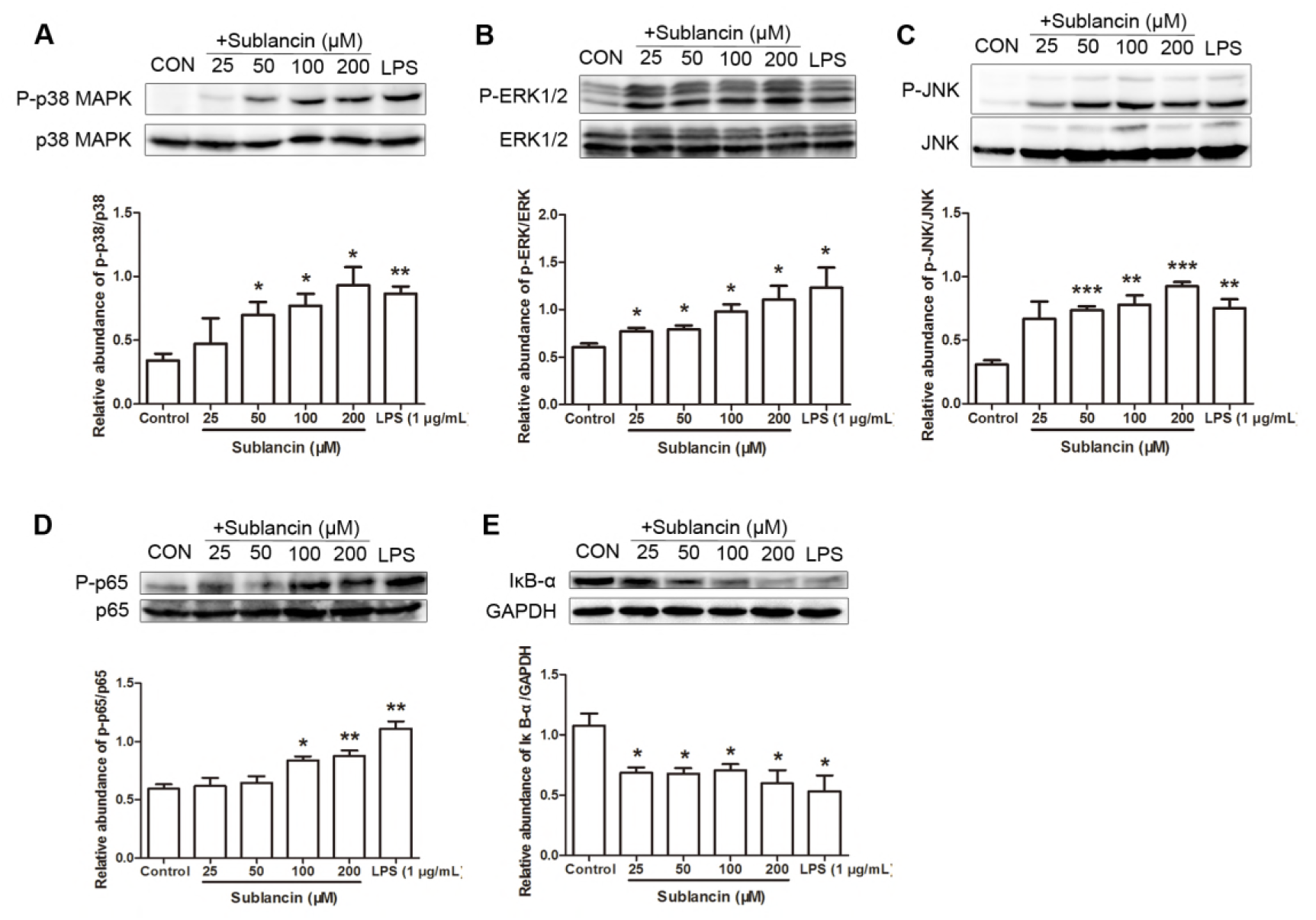
Sublancin regulates the MAPK and NF-κB signaling pathways in macrophages. Representative Western blots for phosphorylated p38 and total p38 (A), ERK1/2 (B), JNK (C) and p65 (D) in RAW264.7 cells treated with sublancin (0-200 μM) or LPS (1 μg/mL) for 30 min. Relavite abundance was represented as phosphorylated protein to total protein expression. (E) Western blot analysis and quantification of IκB in RAW264.7 cells treated with sublancin (0-200 μM) or LPS (1 μg/mL) for 30 min. The values are presented as mean ± SEM (n = 3). Significant differences with control cells were designated as \**P* < 0.05, \*\**P* < 0.01 or \*\*\**P* < 0.001

### *In vivo* sublancin-mediated immune modulation

To evaluate the *in vivo* effect of sublancin, mice were orally administered with sublancin for the indicated time, and P-Mac were collected to assess phagocytosis activity and the capability of TNF-α production. Compared to the control group, the three levels of sublancin treatments (1.0 mg/kg for 28 days; 0.6 mg/kg and 1.2 mg/kg for 14 days) significantly enhanced the phagocytosis activity of P-Mac (Fig. 7A and B), which is consistent with the data of the phagocytosis activity obtained from the *in vitro* study. However, no significant TNF-α promotion was found in P-Mac isolated from sublancin-treated mice (Fig. 7C). It has been reported that NK cells play a central role in the innate immune response to tumors and infections (28). The sublancin treatment of 1.2 mg/kg had a tendency (*P* = 0.0713) to promote NK cell activity compared with the control group (Fig. 7D). Activated T cells (anti-CD3 and anti-CD28 stimulation for 24 hours) from the spleen were examined for IFN-γ production. As shown in Fig. 7E, spleen T cells from 1.2 mg/kg sublancin treated mice secreted more IFN-γ than that from control (*P* < 0.05).

**FIG 7.**
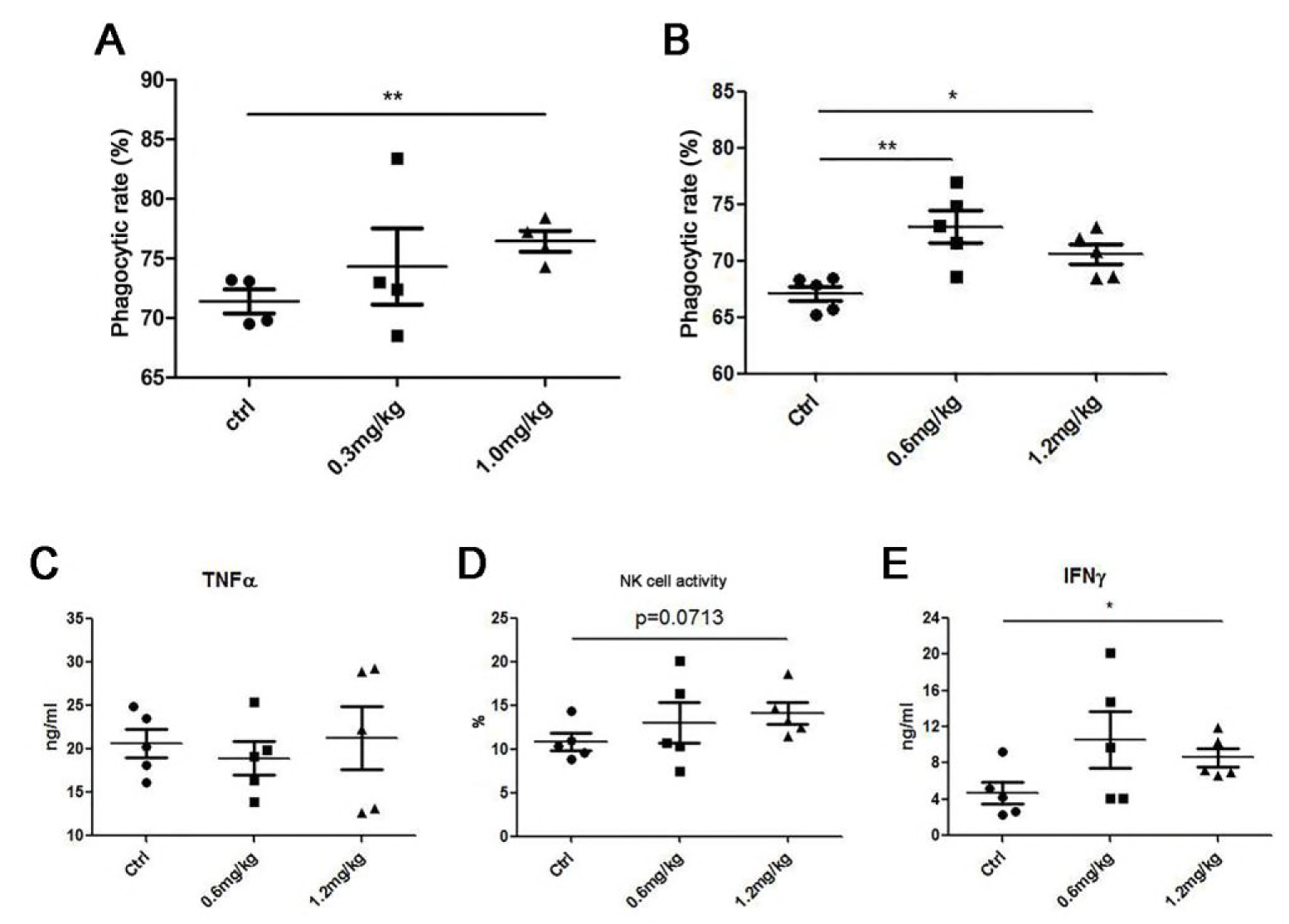
Effects of sublancin on the function of peritoneal cells and splenocytes *in vivo*. Sublancin enhanced the phagocytic activity of peritoneal macrophages *ex vivo* (A and B). Four to six weeks old female BALB/c mice were separated into three groups– control, sublancin low-dose group (0.3 mg/kg or 0.6 mg/kg), and sublancin high-dose group (1.0 mg/kg or 1.2 mg/kg). The mice were orally administered with sublancin for 28 days (0.3 mg/kg and 1.0 mg/kg) or 14 days (0.6 mg/kg and 1.2 mg/kg). Mice in the control group were given by gavage with distillated water. Peritoneal macrophages were harvested and incubated with fluorescent microspheres for determination of phagocytic activity. (C) Comparison of TNF-α production in peritoneal macrophages stimulated with LPS (1 μg/mL). (D) Sublancin (1.2 mg/kg) promoted a weak level of NK cell activity. Splenocytes were isolated from mice and co-cultured with Yac-1 cells by the ratio (Splencytes: Yac-1) of 75:1 for 4 h. Cytotoxicity was measured by the lactate dehydrogenase assay. (E) Comparison of IFN-γ level in activated splenic T cells (2 μg/mL anti-CD3 and 1 μg/mL anti-CD28 antibodies). Data shown are means ± SEM and derived from 4 to 5 mice in each group. Significant difference with control group were designated as \**P* < 0.05, \*\**P* < 0.01

### Protection of mice from MRSA-induced sublethal infection by sublancin

Because of the potent and promising immunomodulatory property of sublancin, this peptide was tested for its anti-infective potential. Sublancin was administered by gavage daily at the indicated doses for 14 consecutive days. Twenty-four hours after the last drug administration, mice were subjected to a sublethal dose (1.0 × 10^7^ CFU/mouse) of MRSA ATCC43300. As shown in Fig. 8, treatment with the three sublancin levels (0.6, 1.0, and 1.2 mg/kg) tended to decrease (*P* < 0.1) the number of viable bacterial counts in the peritoneal fluid 72 hours after infection. Additionally, mice treated with 1.2 mg/kg sublancin had fewer (*P* < 0.05) viable bacterial counts in the peritoneal lavage than the control mice 36 hours after infection.

**FIG 8.**
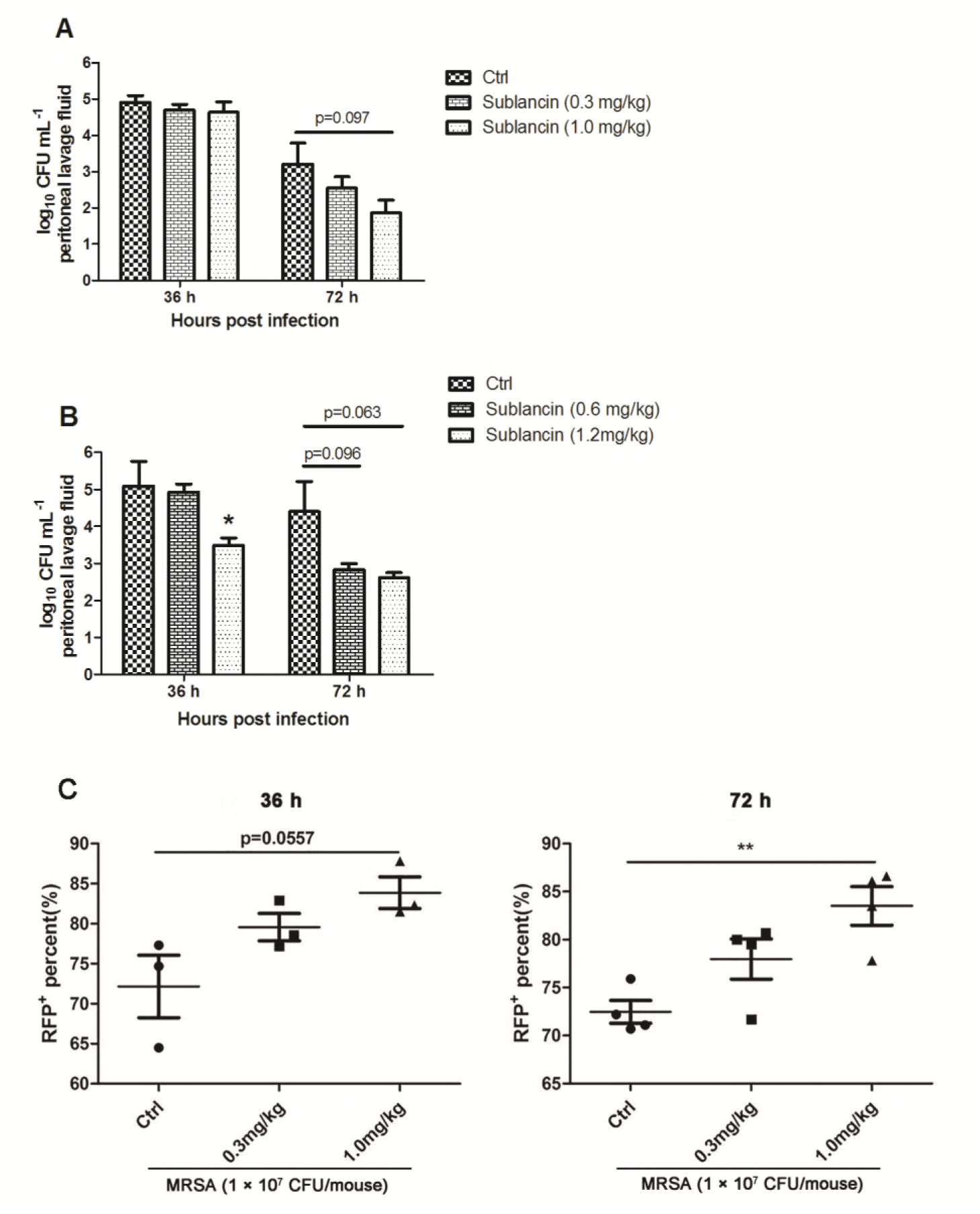
Efficacy of sublancin in MRSA-induced sublethal infection model. (A) Mice were orally administered with sublancin at 0.3 mg/kg or 1.0 mg/kg once a day for 14 consecutive days before intraperitoneal inoculation of MRSA ATCC43300. The staphylococcal load in the peritoneal lavage was gavage was enumerated at 36 and 72 h of infection. (B) Sublancin (0.6 mg/kg or 1.2 mg/kg) was administered by gavage daily for 14 days before infection. Mice were analyzed for bacterial counts in the peritoneal lavage at 36 and 72 h of infection. (C) Peritoneal macrophages were collected 36 and 72 hours after infection for red fluorescent protein (RFP) labeled *Escherichia coli* phagocytosis assay.Data shown are means ± SEM and derived from 3 to 5 mice in each group. Significant difference with control group were designated as \**P* < 0.05, \*\**P* < 0.01

Macrophages have been shown to phagocytose and directly kill bacteria. Therefore, P-Mac were collected 36 and 72 hours after infection for red fluorescent protein (RFP) labeled *Escherichia coli* phagocytosis assay. Compared with control group, the sublancin (1.0 mg/kg) treatment stimulated the phagocytotic activity of P-Mac (Fig. 8C), which may account for the decreased number of viable bacterial counts in the peritoneal lavage. As T cells, especially CD4^+^ and CD8^+^ subsets of T cells, play a critical role in the immune responses to specific antigenic challenges, the changes of T cells in mesenteric lymph nodes (MLNs) were examined. Compared with control mice, mice treated with 1.0 mg/kg sublancin displayed an dramatic increase in the frequency of CD4^+^ and CD8^+^ T cells, with a corresponding increase in the frequency of naïve CD4^+^ T cells and memory CD8^+^ T cells (Fig. 9). This difference was reflected by increased total number of CD8^+^ T cells, naïve CD4^+^ T cells, as well as increased number of memory CD8^+^ T cells and naïve CD4^+^ T cells (Fig. 9). Similarly, mice treated with 1.2 mg/kg sublancin showed increased frequencies of CD4^+^ (*P* < 0.05) and CD8^+^ (*P* = 0.0644) T cells 36 hours after infection (Fig. 10). The frequencies of activated CD4^+^, naïve CD4^+^ and naïve CD8^+^ T cells were significantly increased in MLNs of 0.6 mg/kg and 1.2 mg/kg sublancin treated mice (*P* < 0.05) (Fig. 10). However, no differences in T cell frequency or cell number were observed in MLNs 72 hours after infection among the different treatments (Fig. S2).

**FIG 9.**
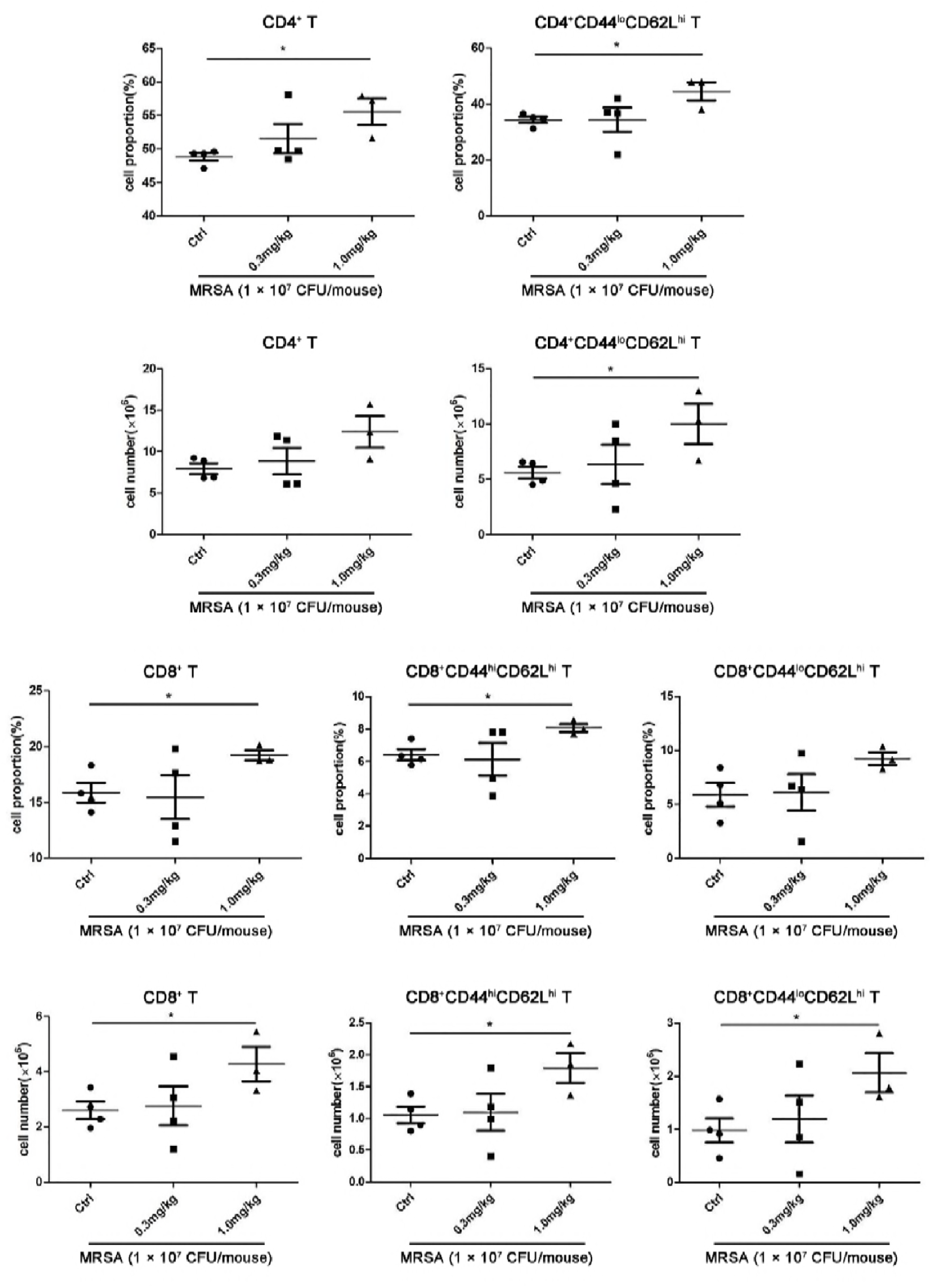
Effects of sublancin on the immune cell subset in the mesenteric lymph nodes (MLNs) of MRSA challenged mice. Mice were orally administered with sublancin (0.3 mg/kg or 1.0 mg/kg) for 14 consecutive days before infection. Phenotypic analysis of single-cell suspensions collected from the MLNs (the MLNs were excised from the mice 36 hours after infection). The phenotype of CD4^+^ and CD8^+^ T cells was further characterized by expression of CD44 and CD62L, cell surface receptors which are differentially expressed on memory (CD44^hi^CD62L^hi^) and naïve (CD44^lo^CD62L^hi^) T cell populations

**FIG 10.**
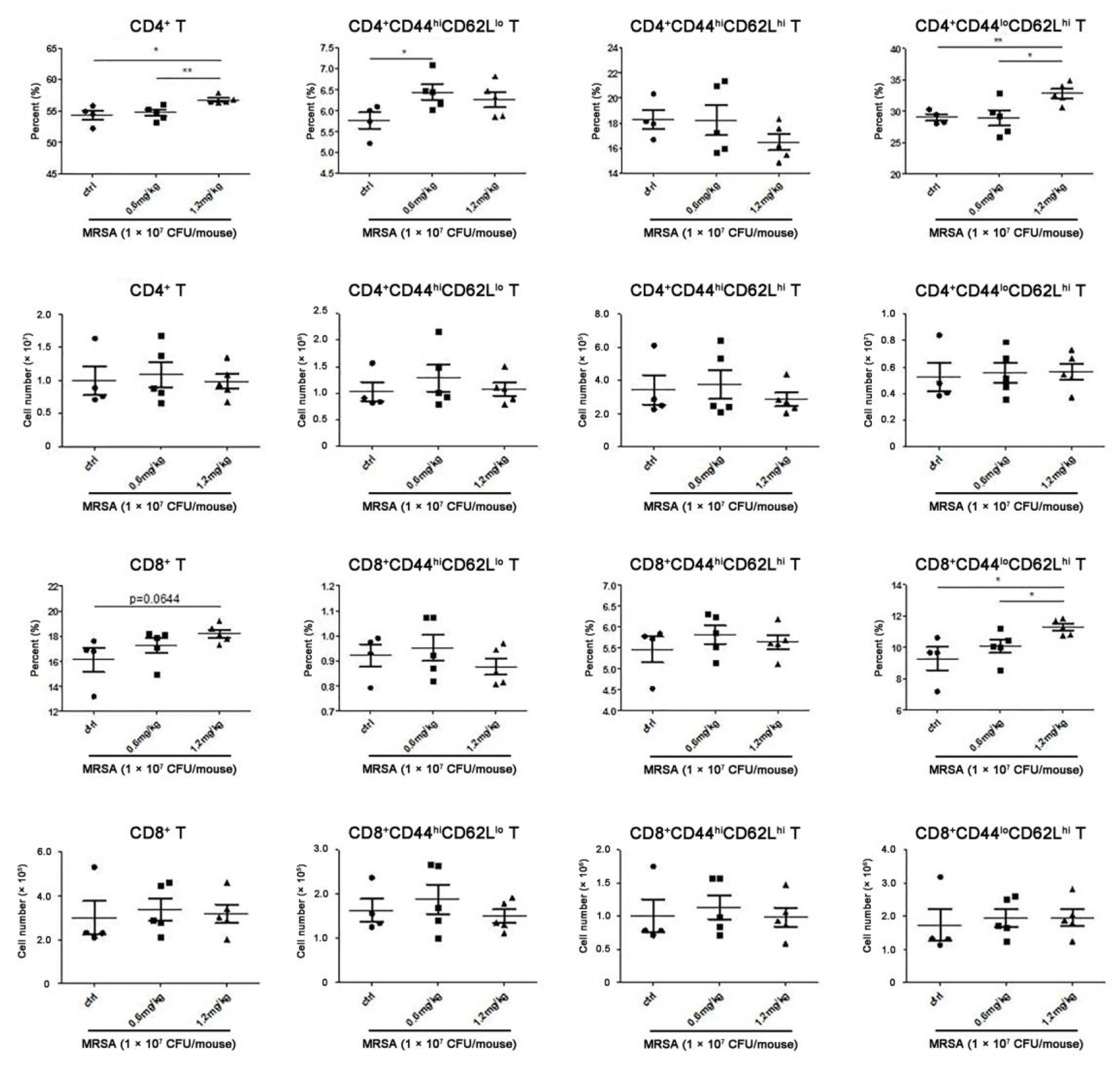
The alteration of various immune cell percentages and cell numbers in the MLNs of MRSA challenged mice. Mice were orally administered with sublancin (0.6 mg/kg or 1.2 mg/kg) for 14 consecutive days before infection. Phenotypic analysis of single-cell suspensions collected from the MLNs (the MLNs were excised from the mice 36 hours after infection). The phenotype of CD4^+^ and CD8^+^ T cells was further characterized by expression of CD44 and CD62L, cell surface receptors which are differentially expressed on activated (CD44^hi^CD62L^lo^), memory (CD44^hi^CD62L^hi^), and naïve (CD44^lo^CD62L^hi^) T cell populations

## DISCUSSION

Antimicrobial peptides provide immediately effective, non-specific defenses against infections through direct bactericidal activity or through indirect modulation of the host defense system by enhancing immune-responsive cells (29). Sublancin, produced by *Bacillus subtilis* 168, has been studied extensively for its antibacterial mechanisms (30, 31). Following our previous studies of the anti-infective efficacy of sublancin in several *in vivo* infection models (16, 17), we examined its role and anti-infection mechanism in a mouse model of MRSA-induced sublethal infection.

Macrophages activation is the key event in innate immunity for host defense against bacterial infections and many immunomodulatory agents activate immune responses primarily by activation of macrophages (32, 33). Activated macrophages are considered to be associated with generation of IL-1β, IL-6, TNF-α and NO. IL-1 is a cytokine that plays pivotal roles in regulating inflammatory responses to sterile insults and infections (34). IL-6 is considered as a key mediator of the acute inflammation (35). TNF-α has many functions such as activation and chemotaxis of leukocytes to kill microbes (36). NO, synthesized by iNOS, is a short-lived gas that possess beneficial roles in antibacterial activity of macrophage against pathogens (37, 38). Upon stimulation by AMPs, different immune cells have been demonstrated to produce cytokines or chemokines. For example, human neutrophil peptides-1 and -3 were found to stimulate the production of IL-1, IL-4, IL-6, and TNF-α in monocytes (39), and MCP-1 in lung epithelial cells (40). In addition, the AMP LL-37 was shown to increase the release of IL-8 in airway epithelial cells (41). Here we show that sublancin stimulated both the protein and mRNA levels of IL-1β, IL-6, TNF-α, and NO in RAW264.7 cells and mouse peritoneal macrophages. Consistently, the mRNA expression of chemokines (such as IL-8 and MCP-1) and costimulatory molecules (B7-1 and B7-2) were also increased by sublancin in RAW264.7 cells and mouse peritoneal macrophages. These observations suggest that sublancin could directly enhance the activation of macrophages.

A critical finding in this study was that sublancin significantly enhanced phagocytic activity of macrophage against MRSA both *in vitro* and *in vivo*. Wan et al. (2014) reported that LL-37 elevated bacterial phagocytosis by human macrophages (42). Furthermore, it has also been demonstrated that human neutrophil peptides 1–3 enhance bacterial phagocytosis by macrophages (43). Consistent with the above notions, we found that sublancin can simulate phagocytosis of fluorescent microspheres in RAW264.7 cells and peritoneal macrophages. Next, the effect of sublancin on the phagocytic activity was determined using mouse peritoneal macrophages *ex vivo*. In healthy or MRSA challenged mice, the oral administration of sublancin enhanced the phagocytotic activity of peritoneal macrophages. Mice that pre-treated with sublancin before the MRSA infection acquired a potent antibacterial activity. It was found that sublancin tended to decrease the bacterial burden in the peritoneal cavity. Our previous research has shown that monocytes/macrophages are pivotal for the protective effect of sublancin (18). Therefore, we further tested whether sublancin could enchance macrophage function by augmentation of the bacterial clearance. In the present study, we found that preincubation of macrophages with sublancin promoted the MRSA-killing activity in macrophages. We speculate that sublancin-enhanced microbicidal activity of macrophages may be due to the the activation of macrophages induced by sublancin. Futher studies are clearly warranted to elucidate the property of sublancin in facilitating bacterial clearance by macrophages.

Increasing evidence has demonstrated that the signaling via TLRs leads to production of a mass of proinflammatory mediators including cytokines (such as IL-1, IL-6, and TNF-α) and NO (44). TLR4 is known to stimulate the innate immune response through recognizing specific molecular patterns. The AMP murine β-defensin 2 has been shown to act directly on dendritic cells through TLR4 (45). In the study presented here, sublancin-upregulated mRNA expression of IL-1β, IL-6, TNF-α, and iNOS was drastically suppressed by TLR4 (TAK-242) inhibitor, suggesting that sublancin activates macrophages probably via the TLR4 signaling pathway. NF-κB and MAPKs are key transcription factors activated in TLR signaling (46, 47). It is well known that NF-κB and MAPKs are involved in regulating cytokine release via phosphorylation of transcription factors. We found that sublancin induced phosphorylation of the three MAPKs (p38 MAPK, ERK1/2, and JNK) in RAW264.7 macrophages. In addition, both NF-κB p65 and IκB-α were responsive to sublancin stimulation by enhancing phosphorylation of NF-κB p65 and the IκB-α degradation, indicating that NF-κB and MAPK signaling pathways were responsible for sublancin-induced macrophage activation. Furthermore, sublancin-induced mRNA expression of iNOS was significantly inhibited by ERK (U0126) and p38 (SB203580) inhibitors. The NF-κB (Bay11-7082) inhibitor also decreased the mRNA expression of IL-1β and IL-6 in RAW264.7 macrophages stimulated with sublancin. These observations highlight the prominent role for the ERK1/2, p38 and NF-κB pathways in the sublancin-induced macrophage activation. Additionally, sublancin-upregulated phosphorylation of p38 MAPK, ERK1/2, and JNK was obviously depressed by TLR4 (TAK-242) inhibitor, further confirming that sublancin probably acts directly on macrophages through TLR4 signaling pathways.

The results obtained from the *in vitro* assays prompted us to explore the *in vivo* immunomodulatory proterties of sublancin in mice. Firstly, the peripheral hemogram analysis from mice that were treated with sublancin (0.3 mg/kg and 1.0 mg/kg) for 28 days showed no disturbances in the blood parameters (data no shown), implying sublancin did not cause apparent toxicity *in vivo*. Moreover, we did not detect any significant differences in total number of splenic cells and peritoneal cells, as well as immune cell subset in the spleen (CD4^+^ T cell, CD8^+^ T cell, B cell, and NK cell) and peritoneal (peritoneal myeloid cell, B cell, macrophage, and neutrophil) cells of mice under healthy conditions (Wang et al., accepted for publication). These findings suggest that sublancin does not alter leukocyte distribution in mice under healthy conditions. IFN-γ has been recognized as the pivotal cytokine of Th1-polarized immunity (48). In the study presented here, sublancin enhanced the ability of spleen T cells to secrete INF-γ, indicating that sublancin slightly triggered the Th1 immunity response.

Importantly, treatment with sublancin protected mice from sublethal infection induced by MRSA ATCC43300. This protection was demonstrated as a siginificantly accelerated clearance of bacteria and mediated partly by enhanced macrophage function. Additionally, emerging evidence also shows that T cell-mediated immunity is crucial to protect against *S. aureus* infection (49). We present here that sublancin stimulated a dramatic increase in the frequencies of CD4^+^ and CD8^+^ T cells in MLNs 36 h postinfection. It is considered that retention of naïve CD4^+^ and CD8^+^ T cells reflects a better immune response (50, 51). In addition, it was also reported that cellular memory responses are also critical for the anti-staphylococcal immunity (52). In the study presented here, sublancin increased the frequencies of naïve CD4^+^ and CD8^+^ T cells, as well as memory CD8^+^ T cells in MLNs at 36 hours after infection. However, there was no siginificant difference in T cell frequency in MLNs among the different treatments at 72 hours after infection. These findings imply that sublancin could improve T cell-mediated immunity in an early infection phase.

In summary, sublancin might enhance not only T cell-mediated immunity but also macrophage function. We posit that sublancin is an excellent alternative to counter the MRSA infections and is worthy of further investigation to successfully translate sublancin to the clinic.

## MATERIALS AND METHODS

### Mice, cell lines, peritoneal macrophages and chemicals

Female BALB/c mice were used for the experiments. The murine macrophage cell line RAW264.7 was obtained from China Infrastructure of Cell Line Resource (Beijing, China) and maintained in Dulbecco’s Modified Eagle’s Medium (DMEM) containing 10% fetal bovine serum (Life Technologies). Peritoneal macrophages (P-Mac) were isolated from BALB/c mice as previously described (53). Briefly, mice were intraperitoneally injected with 2 ml 4% thioglycollate. Three days after injection, peritoneal exudate cells were harvested by lavaging the peritoneal cavity with sterile ice-cold Hank’s balanced salt solution (HBSS). These cells were incubated for 2 h, and adherent cells were used as peritoneal macrophages. Sublancin was generated in our laboratory using a highly efficient expression system involving *Bacillus subtilis* 800 as described privously (17). The purity of this peptide was above 99.6% as determined by high-performance liquid chromatography. Sublancin was produced as lyophilized powder and the endotoxin concentration of the peptide was less than 0.05 EU/mg, as detected by E-Tocate kit (Sigma). Sublancin was resuspended in endotoxin-free water (Sigma-Aldrich) and stored at -20°C. All reagent used in this study were tested for endotoxin to eliminate the interference of endotoxin contamination.

### Cytokine assays

The culture supernatants of RAW264.7 cells or mouse peritoneal macrophages treated with sublancin (25, 50, 100, or 200 μM) for 24 h were collected for the detection of IL-1β, IL-6, and TNF-α levels using commercially available cytometric bead arrays (BD Biosciences) according to the protocol of the manufacturer. Data were acquired with a FACSCalibur flow cytometer and analyzed with BD CBA Software (BD Biosciences).

### NO production

The nitrite accumulated in the culture medium was determined by Griess reaction. RAW264.7 cells or mouse peritoneal macrophages were treated with various concentrations of sublancin (25, 50, 100, or 200 μM). After 24 h, culture supernatants were mixed with an equal volume of Griess reagent (1% sulfanilamide, 0.1% *N*-1-naphthylenediamine dihydrochloride in 2.5% phosphoric acid) and incubated at room temperature for 10 min. The absorbance was detected at 540 nm and NO concentration was determined from a calibration curve of standard sodium nitrite concentrations against absorbance.

### Quantitative real-time PCR

To detect the effect of sublancin on gene expression, RAW264.7 cells or mouse peritoneal macrophages (1 × 10^6^ cells/well) were preincubated on 6-well plates and were treated with sublancin (25, 50, 100, or 200 μM) for 12 h at 37 ^o^C in an atmosphere containing 5% CO_2_. Total RNA were isolated using a RNeasy kit (Qiangen, Hilden, Germany). The quality and quantity of total RNA were determined by gel electrophoresis and a NanoDrop Spectrophotometer (P330, Implen, Germany). cDNA was synthesized from the extracted RNA (1 μg) using a PrimeScript 1st Strand cDNA Synthesis Kit (Takara, Ostu, Japan) according to the manufacture’s protocol. Real-time PCR was performed on an Applied Biosystems 7500 Real-Time PCR System (Applied Biosystems, Singapore) with SYBR Green PCR Master Mix (Takara, Ostu, Japan). Relative gene expression data were normalized against GAPDH and analyzed using the 2^−ΔΔCt^ method (54). Primers for the selected genes are given in Table I.

**Table 1.**
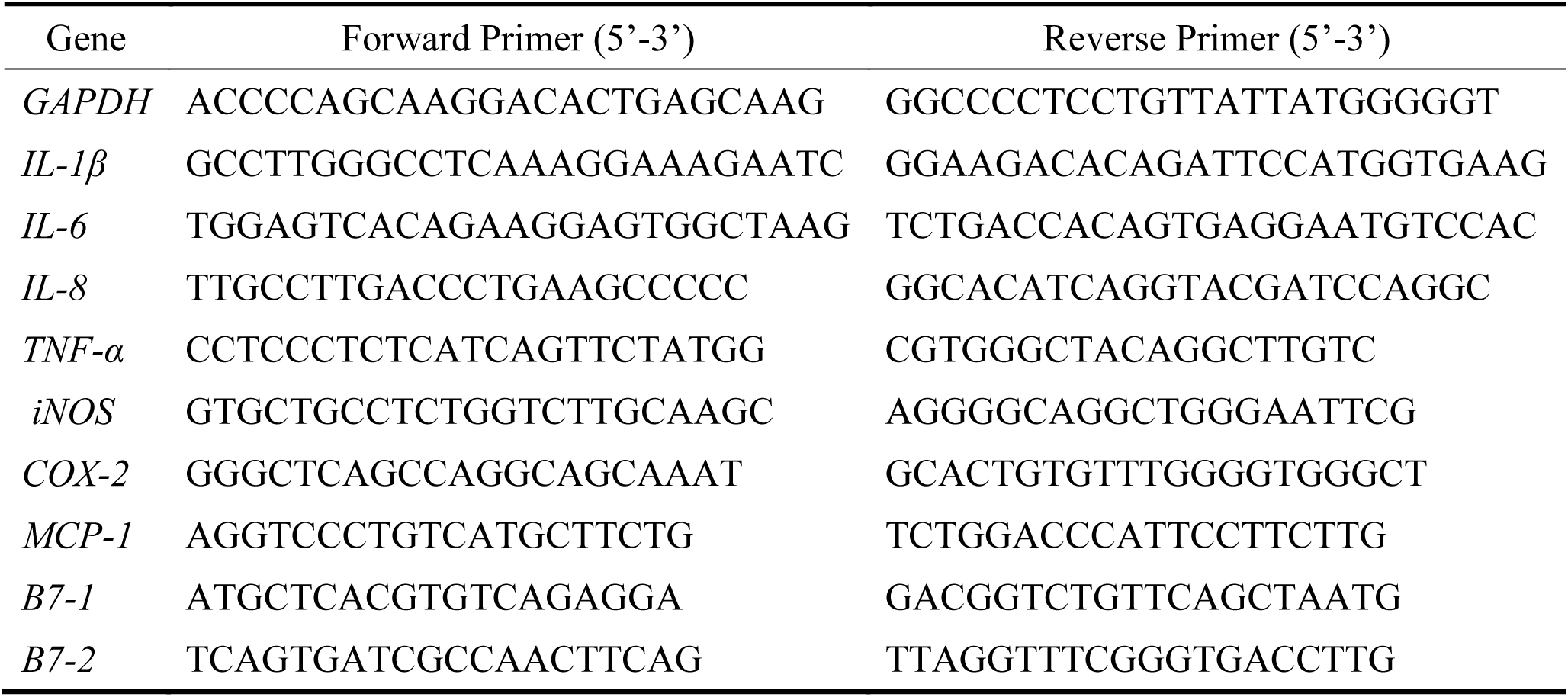
Sequence pf the primers used for quantitative PCR

### Measurement of phagocytic uptake

RAW264.7 cells (1 × 10^6^ cells/well) or mouse peritoneal macrophages were cultured in 6-well plates until 80% confluent. The cells were treated with various concentrations of sublancin (25, 50, 100, or 200 μM) for 12 h. Thereafter, 100 μL of suspended fluorescent microspheres in PBS was added to the wells (cells to beads ratio 1:20) and the cells were incubated at 37°C for 1 h. Phagocytosis was terminated by the addition of 2 mL of ice-cold PBS, and then the cells were washed three times with cold PBS and harvested. Flow cytometric analysis was performed using a FACS Calibur flow cytometer using CellQuest software (BD Biosciences, San Jose, CA, USA).

### Determination of macrophage MRSA-killing activity

MRSA ATCC43300 were grown overnight at 37°C in LB broth, washed in PBS, and adjusted to 10^7^ CFU/ml in DMEM medium. RAW264.7 cells or mouse peritoneal macrophages (2 × 10^5^ cells/well) in 24-well plates were treated with or without sublancin (25 μM) for 12 hours. After being washed with antibiotic-free DMEM medium, cells were incubated with MRSA ATCC43300 for 1 hour (20 bacteria/macrophage). After infection, nonadherent bacteria were washed away using PBS, and macrophages were incubated for 30 minutes (for phagocytosis) or 24 hours (for bacteria killing) in the presence of 10 μg/ml lysostaphin to eliminate the remaining extracellular bacteria. Intracellular bacteria were released by lysing the macrophages in 0.1% Triton X-100, and the number was determined by plating serial dilutions of cell lysates on agar plates. The bacterial killing was expressed as percent changes in bacteria counts using the formular: (bacterial count at 24 hours/bacterial count at 1 hour) × 100.

### Western blot analysis

The RAW264.7 cells grown in 100-mm dish were treated with 100 μM sublancin for the indicated time periods, or sublancin (25 μM, 50 μM, 100 μM, 200μM) for 30 min. Protein was extracted by incubating the RAW264.7 cells with ice-cold lysis buffer containing 150 mM NaCl, 1% Triton X-100, 0.5% sodium deoxycholate, 0.1% SDS, 50 mM Tris-HCl at pH 7.4, and a protease-inhibitor cocktail (Apply Gene, Beijing, China) for 30 min. Subsequently cell extracts were centrifuged at 12,000 × g on 4°C for 10 min. Protein containing supernatant was collected and quantified using a BCA Protein Assay Kit (Pierce, Rockford, IL, USA). Fifty μg of protein samples were electrophoresed on SDS polyacrylamine gels and electrotransferred to polyvinylidene difluoride membranes (Millipore, Bedford, MA, USA). Membranes were blocked with 1 × TBST containing 5% of BSA (Sigma-Aldrich, St Louis, MO) for 2 h at room temperature. The membranes were incubated with corresponding primary antibodies (1:1000 dilution for overnight at 4°C) against p-p38 (Thr180/Tyr182) (Cell Signaling Technology, Cat: 4511S), p38 (Cell Signaling Technology, Cat: 8690S), p-ERK1/2 (Thr202/Tyr204) (Cell Signaling Technology, Cat: 4370S), ERK1/2 (Cell Signaling Technology, Cat: 4695S), p-JNK (Thr183/Tyr185) (Cell Signaling Technology, Cat: 4668S), JNK (Cell Signaling Technology, Cat: 9252S), p-NF-κB (Ser536) (Cell Signaling Technology, Cat: 3033P), NF-κB (Cell Signaling Technology, Cat: 8242P), IκBα (Cell Signaling Technology, Cat: 4812S), and GAPDH (Santa Cruz Biotechnology, Cat: sc-25778). After washing of membranes with 1 × TBST membranes were incubated with a secondary antibody (horseradish peroxidase-conjugated goat anti-rabbit IgG) (Huaxingbio Biotechnology, Beijing, China, Cat: HX2031) at a ratio of 1:10,000 dilution for 1 h at room temperature. The chemifluorescene was detected with the Western Blot Luminance Reagent (Applygene, Beijing, China) using an ImageQuant LAS 4000 mini system (GE Healthcare), and quantified using a gel-imaging system with Image Quant TL software (GE Healthcare).

### Animal experiments

Femele BALB/c mice of 4–6 weeks old were used for all studies. Mice were obtained from HFK Bioscience Co., Ltd. (Beijing, China). All mice used in this study were housed in plastic cages under 12 h light/dark cycle and had access to food and water *ad libitum*. All the techniques, care, and handing of the animals were approved by the China Agricultural University Institutional Animal Care and Use Committee (ID: SKLAB-B-2010-003).

### Sublancin-mediated immune modulation in mice under healthy conditions

The mice were randomly divided into three groups: control, sublancin low-dose group (0.3 mg/kg or 0.6 mg/kg), and sublancin high-dose group (1.0 mg/kg or 1.2 mg/kg). Mice were given sublancin by gavage once a day for 28 days (0.3 mg/kg and 1.0 mg/kg) or 14 days (0.6 mg/kg and 1.2 mg/kg). The control group was orally administered with distillated water daily. Twenty-four hours after the last dose, the animals were killed, and blood was withdrawn for peripheral hemogram analysis. Under an aseptic technique, a laparotomy was performed through a midline incision, and peritoneal macrophages were harvested for phagocytosis assay. The peritoneal cells and spleen were collected for culture.

### *In vivo* efficacy against MRSA-induced sublethal infection

To study the anti-infective role of sublancin in an experimental model of MRSA-induced sublethal infection, mice were randomly allocated to one of three groups: (i) untreated, (ii) sublancin low-dose group (0.3 mg/kg or 0.6 mg/kg), and (iii) sublancin high-dose group (1.0 mg/kg or 1.2 mg/kg). Mice were orally administered with sublancin at the indicated doses for 14 consecutive days, while the untreated mice were given distillated water during the same periods. Twenty-four hours after the last drug administration, all mice were given a sublethal dose of MRSA ATCC43300 (1.0 × 10^7^ CFU/mouse) intraperitoneally. Animals were euthanized 36 and 72 hours after infection and peritoneal lavage was collected in 2 mL of cold HBSS. The staphylococcal load in the peritoneal lavage was enumerated as described previously (55). Peritoneal macrophages were collected for RFP labeled *E. coli* phagocytosis assay. The MLNs were excised for flow cytometry.

### NK cell activity

Splenocytes were collected from mice and co-cultured with Yac-1 cells to obtain an E: T (Splencytes: Yac-1) ratio of 75: 1 in V-bottomed 96-well plates. After 4 h, cytotoxicity was determined by the lactate dehydrogenase (LDH) assay using a LDH Cytotoxicity Assay Kit (Roche, Basel, Switzerland).

### Cell culture

Single-cell suspensions from spleens were prepared passing cells through a 100 μM strainer. The splenocytes were plated at a density of 1 × 10^6^/mL and stimulated with 2 μg/mL anti-CD3 and 1 μg/mL anti-CD28 antibodies (BD PharMingen, San Diego, CA, USA). The peritoneal macrophages were cultured in the presence of 1 μg/mL LPS in DMEM at a density of 1 × 10^6^/mL. The cells were cultured at 37°C for 24 hours before supernatant collection. Supernatants from splenocytes cultures were collected and analyzed by ELISA for INF-γ. The supernatants from peritoneal macrophages cultures were analyzed for TNF-α. All ELISA kits were purchased from eBioscience (San Diego, CA, USA).

### Flow cytometry

Cells from MLNs were stained with CD62L-PE, CD44-APC, CD4-PerCP-Cy5.5, and CD8-PE-Cy7 at 4°C for 30 minutes and then analyzed by flow cytometry (Gallios; Beckman Coulter, Brea, CA, USA). The antibodies were purchased from BioLegend (San Diego, CA, USA), QuantoBio (Beijing, China), BD PharMingen, or eBioscience.

### Statistical analysis

Data are expressed as means ± SEM and were statistically analyzed using one-way analysis of variance followed by Tukey’s post-hoc test (Prism software, version 5). *P* values < 0.05 were taken to indicate statistical significance.

## ACKNOWLEDGMENT

This work was supported by the National Key Research and Development Program of China (2016YFD0501308) and the Special Fund for Agroscientific Research in the Public Interest (201403047).

